# Nascent protein retention at polysomes reduces kinetic barriers to self-assembly

**DOI:** 10.64898/2026.05.17.725791

**Authors:** Shriram Venkatesan, Alex Von Schulze, Jeffrey J. Lange, Jacob Jensen, Lexie E. Berkowicz, Dai Tsuchiya, Ning Zhang, Jennifer Gardner, Scott McCroskey, Ying Zhang, Selene Swanson, Yan Hao, Malcolm Cook, Tong Shu, Liam J. Holt, Laurence Florens, Jay R. Unruh, Randal Halfmann

## Abstract

Living proteomes are necessarily far from equilibrium. It is paradoxical, then, that reducing the translation of new proteins -- which should promote equilibration -- instead prolongs life. We investigated the impact of translational flux to nucleation barriers that preserve the solubility of proteins destined to form amyloids or other assemblies. By manipulating translation initiation rates directly or indirectly, across yeast and human cells, and across a variety of supersaturable proteins, we find that accelerating translation initiation broadly accelerates nucleation irrespective of their global concentrations. We showed that this effect was confined to polysomes and was enhanced by N-terminal placement or other features that retained the nascent aggregating domain at polysomes. Finally, we show that intrinsically disordered regions with high tendencies to self-associate are specifically positioned to do so co-translationally, providing evidence that cotranslational nucleation has shaped proteome evolution.

## Introduction

Life requires a soluble proteome. Solubility is an inherently transient state for many proteins, however. With time, they accumulate into ordered aggregates known as amyloids. The inexorable march of this process is a defining feature of age-associated diseases and perhaps aging itself (Ciryam et al., 2013; Cuanalo-Contreras et al., 2022; David et al., 2010).

The driving force for this progression is supersaturation-- a nonequilibrium state of proteins whose soluble concentrations exceed the minimum for amyloid formation, yet persist due to the dependence of amyloid formation on a pre-existing template that does not readily form spontaneously. This creates a kinetic barrier that allows the soluble proteins to accumulate in the cell. The de novo formation of an amyloid template, known as nucleation (Khan et al., 2018; Vekilov, 2012) can be so infrequent for some proteins, despite extreme supersaturation, that it may never occur in an organism’s lifetime (Serio et al., 2000). Once formed, however, the nucleus templates the deposition of additional molecules of the corresponding protein until equilibrium is reached. Supersaturation drives not only disease but also processes fundamental to life. Certain innate immune signaling proteins use sequence-encoded nucleation barriers to create a perpetually supersaturated state poised for decisive switch-like activation upon pathogen exposure (Rodriguez Gama et al., 2026). How nucleation occurs in vivo is therefore fundamental to our understanding of both life and death.

Nucleation of purified proteins in vitro results from rare fluctuations in density and conformation. The details are still murky, and nucleation theory constantly evolves to accommodate newly discovered complexities (Chatani and Yamamoto, 2018; Martin et al., 2021; Michaels et al., 2023; Šarić et al., 2016; Zhang and Schmit, 2016). The picture is even less clear *in vivo* where aggregation is dynamically influenced by translation, dilution, clearance, and the changing states of other proteins (Hipp et al., 2019; Meisl et al., 2022; Narayanan et al., 2019).

Cells leverage the process of translation to maintain solubility and suppress the fluctuations that trigger nucleation (Waudby et al., 2019). The ribosome plays a major role in this process by constraining nascent chains to lower the entropic penalty of folding (Streit et al., 2024). Co-translational interactions between nascent polypeptides serve as *de facto* kinetic chaperones to prevent off-pathway aggregation (Bertolini et al., 2021; Shiber et al., 2018; Wruck et al., 2025). Co-translational assembly on the same polysome is especially important for homo-oligomers, which comprise nearly one-third of eukaryotic proteins and function across nearly all essential cellular pathways (Lynch, 2012; Mallik et al., 2025). Such interactions also contribute to the formation of translationally active nascent chain/mRNA condensates that govern mRNA localization and protein fate (Chouaib et al., 2020; He et al., 2025a; Safieddine et al., 2026).

The efficiency of translation initiation and elongation both decline with aging, leading to inefficient co-translational complex assembly and aberrant ribosome collisions that promote aberrant nascent chain interactions (Anisimova et al., 2020; Di Fraia et al., 2025; Kim and Pickering, 2023; Stein et al., 2022; Tomuro et al., 2024). That these may fuel the aggregation driving aging is apparent from the fact that decelerating translation -- whether dietarily, pharmacologically, or genetically -- broadly extends lifespan across model organisms (Kim and Pickering, 2023). At the same time, however, reducing the influx of new proteins in the cell necessarily increases the average age of protein molecules and the burden of accumulated damage. Hence, the positive effect of translation deceleration on lifespan is somewhat paradoxical (Anisimova et al., 2018; Kim and Pickering, 2023). How translation impacts the nucleation of supersaturated proteins responsible for disease progression is not yet clear.

Addressing this question is challenged by the fact that nucleation rates and stabilities of protein aggregates are generally anticorrelated (Chakraborty et al., 2023; Khan et al., 2018; Michaels et al., 2023). Resolving this complexity requires a method to quantify both aggregation and protein concentration at single cell resolution across the many thousands of cells required to sample low frequency spontaneous nucleation events. Distributed Amphifluoric FRET (DAmFRET) is a facile method to do so. DAmFRET treats individual living cells as independent femtoliter-volume reaction vessels such that the frequency of query protein aggregation can be measured as a function of its intracellular concentration in genetically identical cells (Khan et al., 2018). By further manipulating protein translation rates, transcript numbers, and turnover in the cell, we can via DAmFRET systematically probe the role of nonequilibrium flux in aggregation across tractable cell cultures (Khan et al., 2018; Venkatesan et al., 2019).

Here we show using DAmFRET and complementary methods that translational flux --irrespective of protein concentration -- is a tunable knob that broadly accelerates protein nucleation *in vivo.* Specifically, we observe that manipulations to polysome size and the residence time of aggregation-susceptible domains at the polysome greatly influence nucleation barriers across a structurally diverse class of supersaturated proteins. This phenomenon proved to be under cellular control and conserved across human cells and *S. cerevisiae*, suggesting that the tempo of protein production is a fundamental mechanism used by translation machinery to maintain the metastable solubility of the proteome.

## Results

### Proteasome inhibition stalls amyloid formation

We began our study with a well-established model for amyloid nucleation -- the amyloid-beta peptide (Aβ42) (Cohen et al., 2013; Michaels et al., 2018; Seuma et al., 2021; Srivastava et al., 2019). Aβ42 serves as an ideal probe for the mechanisms of nucleation in cells because its assembly is highly sensitive to both concentration and conformation. The concentration sensitivity is clinically evident in early-onset Alzheimer’s cases involving trisomy 21 or secretase mutations, while its susceptibility to seeding events demonstrates a high sensitivity to conformational templates (Novo et al., 2018; Rosenberg et al., 2016; Watts et al., 2014). Using DAmFRET, we observed that HEK293T cells expressing mEos3.2-tagged Aβ42 form two distinct populations that overlap across a range of expression levels (**Fig. S1A**). This pronounced bimodality in AmFRET levels reflects a nucleation-limited transition characteristic of amyloid (Khan et al., 2018). In contrast, Aβ40 does not form amyloid and exhibits a single low AmFRET population (**Fig. S1A**).

To gain total control over protein turnover, we first blocked the major pathway for protein degradation -- the proteasome -- by treating Aβ42-expressing HEK293T cells with Bortezomib (BTZ). While we expected to see enhanced amyloid formation, owing to a buildup of misfolded species, we instead saw the opposite -- a dose-dependent decrease in amyloid nucleation (**Fig. 1A**). This proved true whether BTZ was added right after transfection or instead after 24 hours of prior protein expression (**Fig. S1D,E, Methods**). Importantly, by comparing cells of equal expression levels across treatments, the reduced nucleation with BTZ (rightward shift in the fits) occurred irrespective of any effect on the concentration of Aβ42. To ensure that the effect reflected proteasome inhibition rather than a potential off-target activity of BTZ, we repeated the experiment with other proteasome inhibitors MG-132, Lactacystin and Carflizomib. Again, nucleation was inhibited as seen by a decrease in amyloid formation in a representative overlapping expression range (**Fig. S1B**). To account for any potential artifact of the fluorescent protein tag, we next repeated the experiment with 3xFLAG-tagged Aβ42 and quantified amyloid formation by SDD-AGE of equally loaded lysates. Here too, BTZ-treatment greatly reduced amyloid formation (**Fig. S1C**).

**Figure 1:**
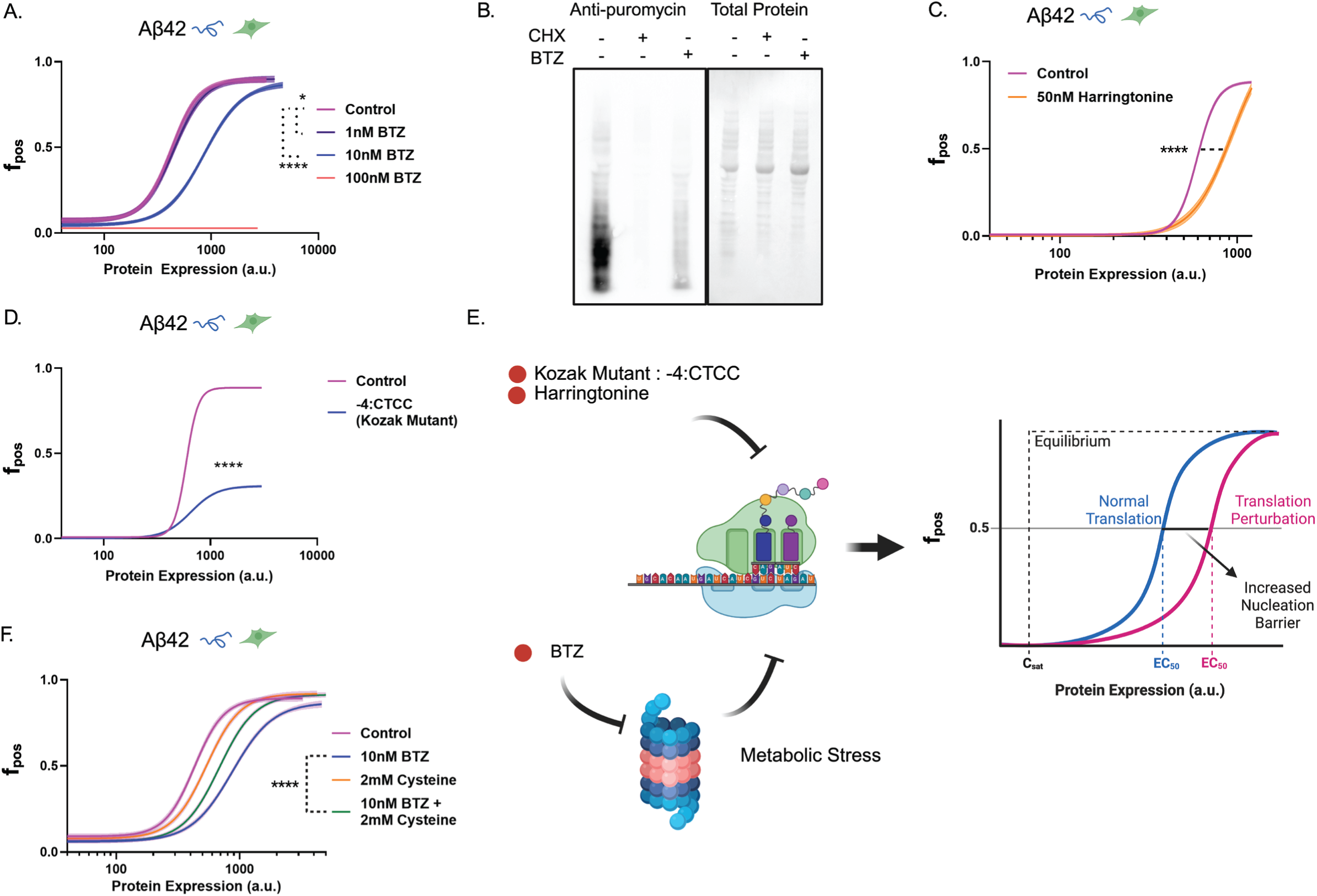
Translational perturbations augment nucleation behavior *in vivo*. (A) DAmFRET analysis reveals that BTZ treatment significantly increases the nucleation barrier of A□42 in a dose-dependent manner (HEK293T cells; 20 hr treatment upon transfection). (B) Protein synthesis rates in HEK293T cells are reduced upon treatment with the proteasome inhibitor Bortezomib (BTZ; 100 nM; 20 hr) as determined by Surface Sensing of Translation (SUnSET). Cycloheximide (CHX) is included as a positive control for translation inhibition. (C–D) Pharmacological inhibition of initiation via Harringtonine (20 hr treatment upon transfection) (C) and targeted genetic suppression of initiation via a weak -4:CTCC Kozak mutation (D) result in drastic reduction in the nucleation of A□42 amyloid. (E) Conceptual model illustrating how global (BTZ, Harringtonine) and transcript-specific (Kozak mutant) perturbations of translation act as a kinetic throttle to regulate amyloid nucleation *in vivo*. Because of the nucleation barrier and the finite time cells are supersaturated prior to analysis, the thermodynamic solubility limit (saturating concentration, C_sat_) is always lower than the observed onset of nucleated polymerization. (F) Supplementation with cysteine (2 mM) provides a significant rescue of the BTZ-induced increase in the A□42 nucleation barrier. (*n=3*; \**p < 0.05*, \*\*\**p < 0.001*, \*\*\*\**p < 0.0001*).

The counterintuitive nature of this result compelled us to investigate the mechanism. We considered whether alternative protein quality control pathways might be overcompensating for the loss of proteasomal degradation under BTZ treatment by degrading or solubilizing protein nuclei before they mature into detectable aggregates. To test for a role of autophagy, we treated cells with Chloroquine (CQ) and Wortmannin (WM). These inhibitors of autophagy had no effect on the BTZ-induced suppression of nucleation (**Fig. S1D**), indicating that enhanced clearance is not responsible for the increased kinetic barrier. We then blocked the primary axis of the heat shock response using a broad spectrum HSP70 inhibitor (VER-155008), but again observed no rescue of the BTZ effect (**Fig. S1E**), suggesting that an upregulation of chaperones induced by proteasome inhibition is also unlikely to explain the reduction in nucleation.

Because amyloid formation requires the repeated arrival of soluble proteins to a growing aggregate, the effect of BTZ could in principle be explained by a global reduction in macromolecular diffusivity that can result from stress (Joyner et al., 2016; Munder et al., 2016; Parry et al., 2014). We used Genetically Encoded Multimeric nanoparticles (GEMs) to evaluate this possibility (Delarue et al., 2018; Hernandez et al., 2024). BTZ treatment caused only a slight (less than 10%) reduction in effective diffusion for 50 nm particles, suggesting that the observed magnitude of suppression of nucleation is unlikely to be attributed to the subtle reduction in mEos-Aβ42 diffusion (**Fig. S1F,G**).

Having ruled out enhanced clearance, chaperone upregulation, and altered diffusivity, we turned to an unbiased proteomic approach to define the system-wide effects of BTZ. We isolated the mEos-Aβ42 interactome from vehicle- and BTZ-treated cells using co-immunoprecipitation (**Fig. S1H**) and analyzed the complex constituents via quantitative mass spectrometry. Pathway enrichment analysis of the proteins significantly downregulated by BTZ revealed a broad reduction in translational machinery (aminoacyl-tRNA biosynthesis, ribosome) and central carbon metabolism (glycolysis, TCA cycle), alongside multiple neurodegenerative disease pathways (**Fig. S1I-K**). This suggests that rather than acting through canonical quality control mechanisms, proteasome inhibition may stall amyloid nucleation by rewiring the metabolic and translational landscape of the cell.

### Translational perturbations modulate nucleation behavior *in vivo*

Proteasome inhibition has been shown to suppress global translation rates by triggering the integrated stress response that phosphorylates the eukaryotic initiation factor 2α (eIF2α) (Fournier et al., 2010; Jiang and Wek, 2005). Non-lethal doses of MG132 and BTZ reduce the rate of degradation while simultaneously triggering a compensatory reduction in the rate of protein synthesis (Bush et al., 1997; Goldberg, 2003), thereby preserving steady state protein concentrations (Bush et al., 1997). Using SUrface SEnsing of Translation (SUnSET) to monitor the rate of protein synthesis through the immunodetection of puromycin-labeled nascent proteins, we confirmed that translation rate indeed declined upon BTZ treatment (**Fig. 1B**).

To probe the effects of translation on nucleation rates more directly, we first perturbed translation initiation pharmacologically. Harringtonine specifically inhibits the first step of peptide bond formation by binding to the A-site of the 60S subunit, thereby stalling ribosomes at initiation codons (Fresno et al., 1977; Ingolia et al., 2011). Similar to the effects observed with BTZ, low-dose treatment with Harringtonine significantly repressed Aβ42 nucleation (**Fig. 1C**).

To isolate the effect of translation to just the query protein, we next mutated the Kozak sequence directly upstream of the Aβ42 ORF, removing the consensus -3 purine to reduce translation initiation efficiency (Noderer et al., 2014). With this suppressive Kozak mutation, we observed a significant decrease in Aβ42 nucleation as revealed by comparison of expression-matched wild-type control (**Fig. 1D**). Critically, this result confirms that suppressing translation initiation alone—without globally changing protein homeostasis or the specific protein’s concentration—suffices to modulate its probability of nucleation in the cell (**Fig. 1E**).

While these results confirm that suppressing translation initiation reduces nucleation, they do not directly establish this as the mechanism by which proteasome inhibition does so. Previous literature suggests that translational repression can be specifically rescued via the supplementation of cysteine, which restores translation despite the continued presence of a proteasome inhibitor (Fresno et al., 1977; Suraweera et al., 2012). Indeed, we observed a significant rescue in BTZ-induced repression Aβ42 amyloid formation upon cysteine supplementation in BTZ-treated cells (**Fig. 1F**). These data confirm that translation rate is a primary modulator of nucleation behavior and align with our proteomics data that metabolic stress is the primary trigger of translational suppression by proteasome inhibition (**Fig. 1E, Fig. S1I-K**).

In summary, we show that acute proteasomal inhibition surprisingly reduces amyloid aggregation through its negative feedback on translation initiation, and that the translation initiation rate of an amyloid-forming protein directly and specifically controls the probability that it will nucleate an amyloid that irreversibly collapses its solubility. Our findings thus far collectively implicate translational control of nucleation as a potential hedge against irreversible losses in solubility under proteotoxic stress.

### Translation governs nucleation across structural classes and biological contexts

While HEK293T cells provide a model for understanding these phenomena in the context of the dynamic human proteome, they cannot address its potential generality across biological contexts. We therefore next asked if translation flux likewise governs nucleation frequencies in a model single celled eukaryote -- budding yeast. DAmFRET in this system is also exceptionally well-established and offers a more systematic and mechanistic dissection of the phenomenon. Namely, nucleation events are quantized by blocking cell division upon query protein expression, while an accurate proxy for cell volume allows protein concentration (instead of total expression levels) to be measured, and phase boundaries to be pinpointed, across an approximately 100-fold range of concentrations (Khan et al., 2018; Kimbrough et al., 2025a; Miller et al., 2023).

Active translation is known to drive stress-induced protein aggregation, presumably through the inherent tendency of nascent polypeptides to misfold (Hartl et al., 2011; Medicherla and Goldberg, 2008; Zhou et al., 2011). To determine if the relationship between translation and nucleation is limited to misfolding-dependent aggregation, we compared two well-characterized and physically distinct classes of self-assemblies: native polymers of globular ɑ-helical death fold domains (DFDs) and □-sheet amyloids formed by intrinsically disordered prion-like domains (PrDs). While these classes differ in their structural architecture, both are characterized by significant kinetic barriers that render nucleation stochastic. DFD polymer nucleation is highly sensitive to the local concentration of DFD protomers, which retain their pre-existing conformation upon polymerization (Rodriguez Gama et al., 2022). PrD amyloid nucleation instead involves a rate-limiting conformational ordering within initially disordered oligomers (Khan et al., 2018; Serio et al., 2000). These disparate classes of self-assemblies are therefore differentially responsive to perturbations in local concentration versus conformational flexibility that could each be impacted by translation rate. Both have been extensively characterized by DAmFRET in yeast cells (Goncharoff et al., 2025; Posey et al., 2021; Rodriguez Gama et al., 2026).

To decrease translation rate while allowing for gradual but continual protein synthesis, we separately utilized nanomolar doses of Rapamycin to inhibit TOR-dependent initiation, Lactimidomycin (LTM) to target the final step of initiation, and Cycloheximide (CHX) to inhibit elongation (Barbet et al., 1996; Schneider-Poetsch et al., 2010). All translation inhibitors reduced nucleation, and did so regardless of the protein assessed (**Fig. 2A**). Therefore, translation initiation broadly governs nucleation frequencies through a conserved physical principle that does not strictly depend on the protein being synthesized.

**Figure 2:**
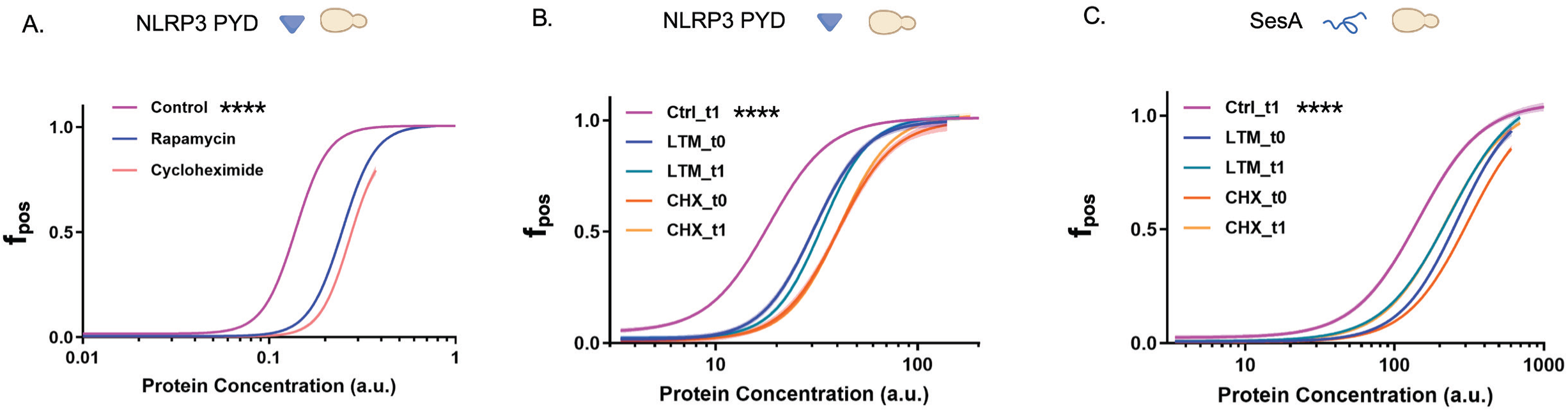
Translational flux is a conserved regulator of nucleation across structural classes. A) Pharmacological inhibition of initiation via Rapamycin (200 nM; 24 hr) or elongation via Cycloheximide (CHX; 180 nM; 24 hr) significantly reduces nucleated-polymerization of the globular death fold domain (DFD), NLRP3 PYD in *S. cerevisiae* (MG132 100 µM). (B–C) Time-course DAmFRET analysis of NLRP3 PYD (B) and the disordered SesA (C) following treatment with CHX (100 µM) or the initiation inhibitor Lactimidomycin (LTM; 3µM) for either 4 hours (t0) or 24 hours (t1). Despite a 24-hour window for stochastic nucleation to occur, translation suppression results in significantly lower nucleation compared to the 24-hour untreated control, identifying that deficiencies in co-translational interaction halt assembly competence. (*n=3*; \**p < 0.05*, \*\*\**p < 0.001*, \*\*\*\**p < 0.0001*).

### Newly synthesized proteins have a transient window for nucleation competence

Because proteins accumulate more rapidly in fast translating cells relative to slow translating cells, the protein molecules in a given bin of total concentration on a DAmFRET plot will be younger on average in fast translating cells. Consequently, the effect of translation rate on nucleation frequency could indicate a preference for nucleation by younger protein molecules irrespective of how they were translated.

To determine if the proteins lose nucleation competence with time, we conducted a simple time course experiment to follow nucleation of pre-supersaturated proteins. For this purpose we used two proteins that tend to nucleate in the mid to upper concentration range accessed in our system -- a globular DFD (NLRP3 PYD (Rodriguez Gama et al., 2026)) and an intrinsically disordered amyloid (SesA (Khan et al., 2018)) -- so as to increase the observation window for new nucleation events. After inducing query protein expression overnight, enough to supersaturate deeply, we treated cells with a high dose of CHX or LTM to stop translation and waited 4 hours (> 10 maturation halftimes (Zhang et al., 2012)) to allow pre-synthesized mEos molecules to mature before taking a baseline DAmFRET measurement. We then measured DAmFRET again after 20 additional hours of treatment. Strikingly, no new nucleation events were detected for NLRP3 PYD and SesA following translation inhibition (**Fig. 2B,C**). Note that in the absence of treatment, cells would only have just reached their final concentration at the time of measurement, while treated cells would have been sitting at that protein concentration for 20 hours. Most nucleation events therefore occurred at lower concentrations than the observed final concentration. That nucleation effectively stopped in the absence of ongoing translation strongly suggests that nucleation only occurs at the time of protein synthesis.

### Nucleation is governed by the proximity of new molecules rather than their age

The above result could reflect strictly time-dependent changes to the molecules, as by post-translational modifications or unbinding of co-translationally loaded chaperones, for example, or it could simply reflect their diffusion away from the site of translation. To dissect these potential contributions, we designed an experiment to perturb initiation rates without changing the age distribution of protein molecules in the cell. To do so, we leveraged the fact that our 2-micron plasmid copy number can be tuned by the strength of selection for the *URA3* auxotrophic marker (Futcher and Cox, 1984; Rose and Broach, 1990). Specifically, we titrated uracil in the medium to relax plasmid selection in cells expressing the strong Kozak construct, while retaining stringent plasmid selection in cells expressing the weak Kozak construct (**Fig. 3A**). We thereby identified a specific condition where at 24 hours of induction cells expressing the strong Kozak construct exhibited a distribution of protein concentrations indistinguishable from cells expressing the weak Kozak construct (**Fig. 3B**). This experimental design created two distinct translational regimes at the same bulk abundance: one with more ribosomes loaded onto fewer mRNAs (strong Kozak/low mRNA) and one with less ribosomes loaded onto more mRNAs (weak Kozak/high mRNA) (Hinnebusch, 2014; Shah et al., 2013). The former regime favored nucleation for both disordered PrDs (Pub1) and structured DFDs (NLRP3 PYD) (**Fig. 3C,D**). The probability of nucleation therefore strongly depends on the proximity of synthesis on the same mRNA. Taken together, these results strongly suggest that the fundamental “unit” of nucleation is the individual polysome.

**Figure 3:**
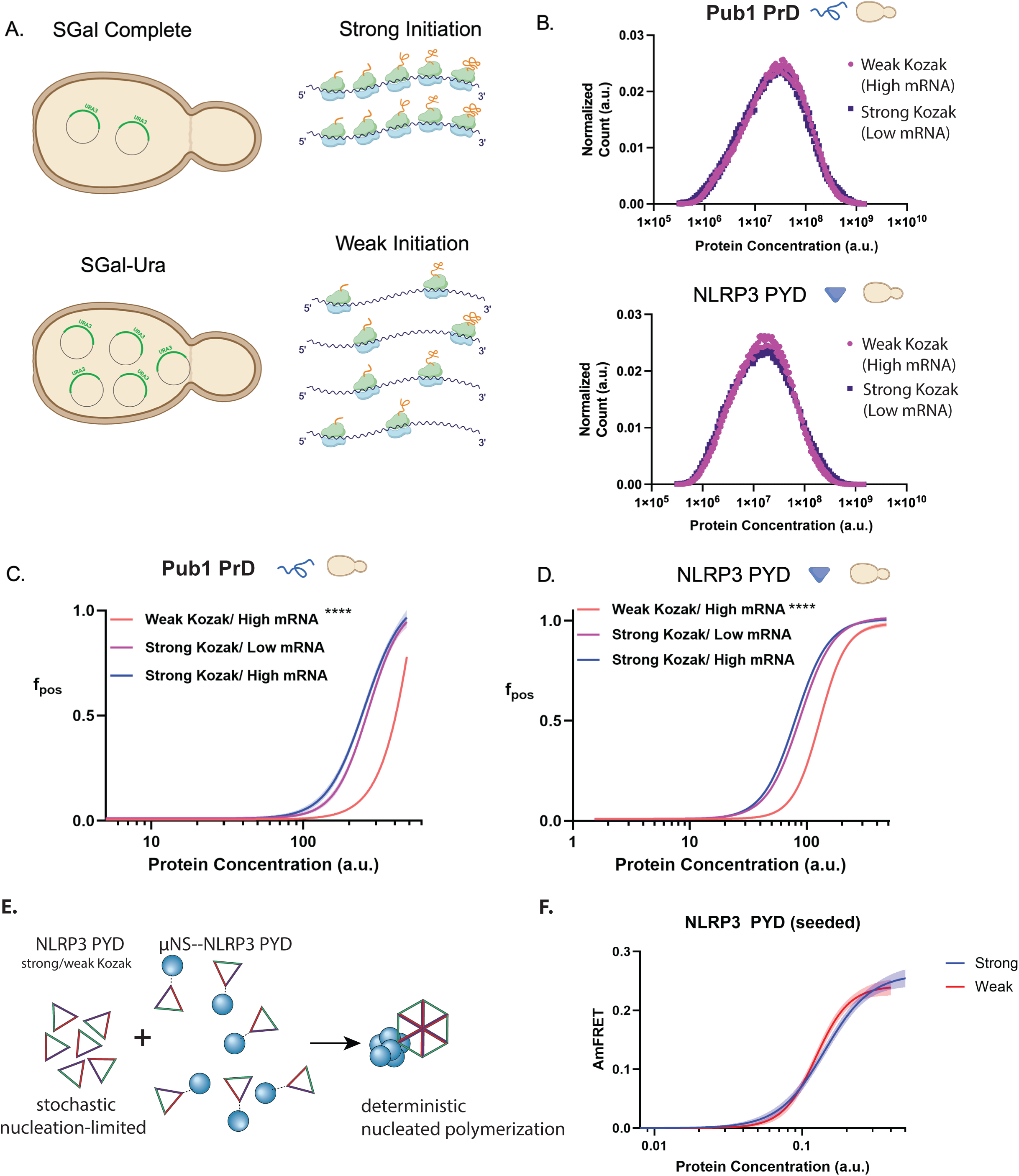
Nucleation is governed by transcript-level ribosome loading rather than global protein abundance. (A) Overview of the Ura-doping strategy in *S. cerevisiae* utilized to isolate the effect of initiation rate from global protein concentration. By titrating uracil levels in the growth medium, we obtained a specific condition where cells expressing a strong Kozak sequence at low plasmid copy numbers exhibited a total protein expression distribution indistinguishable from cells expressing a weak Kozak sequence at high plasmid copy numbers. (B) Representative expression histograms confirming identical endpoint global protein distributions for cells expressing a strong Kozak sequence at low plasmid copy number (SGal complete media) and a weak Kozak sequence at high plasmid copy number (SGal -Ura media) for both Pub1 PrD and NLRP3 PYD, suggestive of equal translation output. (C–D) DAmFRET analysis demonstrating that the increased ribosome loading suffices to increase nucleation for both the Pub1 PrD (*[pin^−^]*) (C) and the NLRP3 PYD (*[PIN^+^]*) (D) despite identical total protein abundance. (*n=3*; \**p < 0.05*, \*\*\**p < 0.001*, \*\*\*\**p < 0.0001*). (E) Schematic of experimental design showing elimination of the nucleation barrier of NLRP3 PYD-mEos expressing artificial NLRP3-PYD condensates in trans. (F) Spline fits of AmFRET trace over the range of protein concentration showing overlapping fits despite Kozak mutation, while allowing NLRP3 PYD polymerization to not be rate-limited by nucleation.

If translation-mediated proximity simply lowers the nucleation barrier (without changing equilibrium solubility), then the self-assembling domain should assemble to the same extent regardless of how it was translated. We directly tested this by fusing NLRP3 PYD to a multivalent viral condensate-forming protein μNS (Rodriguez Gama et al., 2026) and co-expressing these “seeds” in trans with the DAmFRET construct expressing NLRP3 PYD-mEos (**Fig. 3E**). The NLRP3 discontinuity in the unseeded DAmFRET profile was eliminated in the presence of the seeds, implying elimination of the nucleation barrier (**Fig. S2A**). Tracing the spline fit of the profile shows an identical saturating concentration for both strong and weak Kozak variants of NLRP3 PYD-mEos, confirming that translation flux specifically affects the kinetic barrier to self-assembly rather than the thermodynamic stability of the assemblies (**Fig. 3F**).

Further, to ensure that the Kozak-dependence of nucleation was not an artifact of a specific tag (mEos) in the yeast system, we tagged a DFD (ASC PYD) with an orthogonal monomeric fluorogen (frFAST) that is structurally distinct and only half the size of fluorescent proteins. frFAST covalently binds a panel of FRET-compatible, membrane permeable dyes (Li et al., 2020). By titrating equal amounts of two dyes with overlapping emission and excitation spectra -- tfLime and tfPoppy, we obtained an equivalent discontinuous, bimodal “DAmFRET” profile for ASC PYD. Lastly, we observed qualitatively similar dependence of nucleation on Kozak strength using frFAST-tagging as with mEos-tagging (**Fig. S2B**).

### N-terminal placement of the polymerizing domain enhances nucleation

Having established that translation initiation rates govern nucleation through a proximity-dependent mechanism, we next varied the length of time a newly synthesized nucleating domain resides at the polysome. We reasoned that if nucleation is driven by the local density of nascent chains, then the duration and proximity of these chains to one another during synthesis should define the transient window of nucleation competence.

We first tested if the order of ribosomal exit influences nucleation behavior by comparing fusions where the protein of interest (POI) was placed at either the N-terminus or the C-terminus of the mEos tag (**Fig. 4A**). In the N-terminal configuration, the nucleating domain exits the ribosome first but remains tethered while the remainder of the fusion protein is synthesized. In contrast, in the N-terminal configuration, the nucleating domain is synthesized last and released immediately from the ribosome.

**Figure 4:**
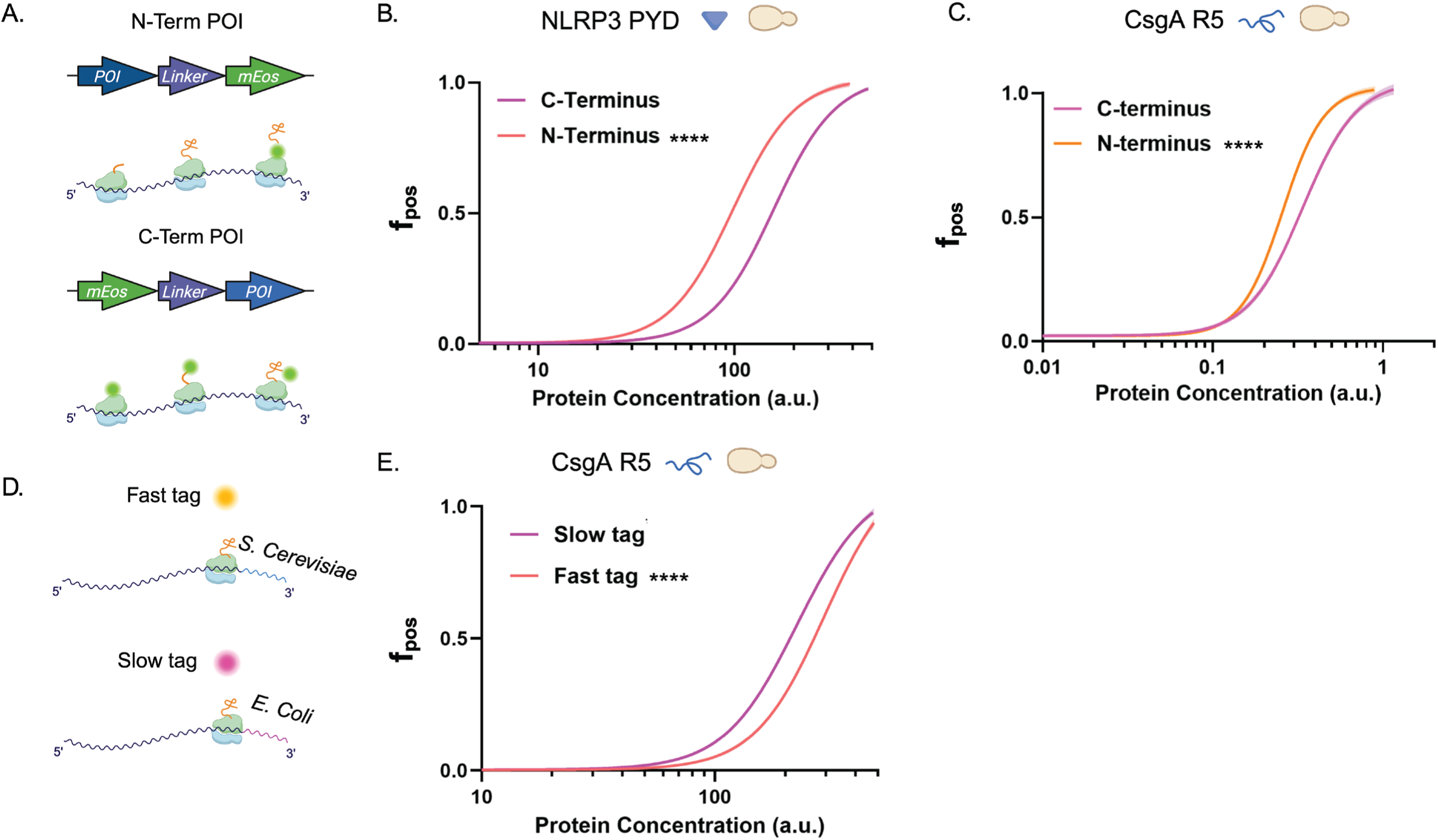
Increased residence time via domain placement or ribosome “pile-up” facilitates assembly. (A) Schematic of constructs designed to test the impact of domain placement on nucleation; the protein of interest (POI) was fused at either the N-terminus or C-terminus of the mEos tag to alter the order of ribosomal exit. (B–C) DAmFRET analysis of the NLRP3 PYD and CsgA R5 reveals that N-terminal placement significantly enhances nucleation, which can be attributed to the C-terminal domain tethering the POI for longer time at the ribosome, allowing it to interact with nascent chains on neighboring ribosomes before diffusing into the cytosol. (D) Strategy to modulate residence time via a downstream ribosome “pile-up”. An innocuous C-terminal tag was encoded using either *S. cerevisiae* optimized codons (fast tag) or *E. coli* optimized codons (slow tag). (E) Evaluation of CsgA R5 nucleation with either tag. Consistent with the residence time hypothesis, the slow *E. coli* tag significantly reduced nucleation to amyloid. (*n=3*; \**p < 0.05*, \*\*\**p < 0.001*, \*\*\*\**p < 0.0001*).

Working initially with NLRP3 PYD, we observed that placing the tag on the C-terminus enhanced self-assembly as expected (**Fig. 4B**). However, given the 100-residue length of NLRP3 PYD relative to the approximately 30-residue length of the ribosome exit tunnel, we reasoned that interactions between nascent NLRP3 PYD polypeptides could confound our interpretation. We therefore repeated this experiment using a very short (19 residue) amyloid forming segment of the *E. coli* Curli protein (CsgA R5 (Wang et al., 2007)) to exclude any possibility that the N-terminally tagged version could aggregate prior to release from the ribosome. Here too, we observed enhanced aggregation when CsgA R5 was retained at the ribosome by the C-terminal tag (**Fig. 4C**). These data suggest that the added residence time provided by cotranslational tethering is a potent driver of aggregation.

### Increased residence time via ribosome “pile-up” facilitates assembly

As an orthogonal test of the effect on nucleation of nascent chain residence time at the polysome, we engineered a system to stall ribosomes downstream of the nucleating domain. To do so, we appended an innocuous intrinsically disordered C-terminal sequence to the amyloid forming protein CsgA R5. We encoded this sequence in either a “Fast Tag” version using *S. cerevisiae-*optimized codons or a “Slow Tag” version using codons optimized for a very different organism (*E. coli*) (**Fig. 4D**). Importantly, the amino acid sequence of the tag did not differ. We reasoned that the Slow Tag would accumulate ribosomes, effectively increasing the duration the upstream nucleating domain remains tethered to the ribosome (Hanson and Coller, 2018). As expected, the Slow Tag greatly increased nucleation even at identical intracellular concentrations (**Fig. 4 E**).

Increased translation initiation and stall-inducing sequences can cause elongating ribosomes to collide, leading to recognition by ribosomal quality control (RQC) machinery and extension of the nascent chains with aggregation-prone C-terminal alanine- and threonine-rich (CAT) tails(Brandman et al., 2012; Shen et al., 2015). To determine if translational flux-driven nucleation depended on this mechanism, we genetically disrupted the RQC machinery. Specifically, we repeated the comparisons between protein pairs with different Kozak sequences or synonymously encoded C-terminal extensions in yeast with the CAT tail installing enzyme Rqc2 (Shen et al., 2015) deleted. If RQC mediates the observed effect of translational flux on nucleation, then loss of Rqc2 should erase the effect of Kozak strength and codon usage in C-terminal extensions. We observed, however, that these variables still dominated nucleation (**Fig. S2C**), indicating that translation flux-driven nucleation cannot be attributed to CAT tail-mediated aggregation, and is therefore presumed to be driven by direct cotranslational association of the aggregating domains at the polysome.

### Endogenous self-assembling IDRs are specifically poised to assemble co-translationally

Nucleation barriers involve phase transitions, or self-assembly beyond finite oligomerization or strictly one-dimensional polymerization. To investigate if endogenous phase transitions may also be under kinetic control by polysomes, we took a bioinformatic approach focused on the largest category of such proteins -- IDRs, for which computational tools can now reliably predict self-association. We specifically analyzed human proteins with annotated IDRs longer than 100 residues whose potential for self-interaction had been evaluated by FINCHES, a computational framework that uses chemical physics to predict IDR-IDR attraction or repulsion (Ginell et al., 2025). We then extracted the predicted translation efficiency (TE) of these ORFs using RiboNN, a recent deep learning model trained on Ribo-seq datasets from 78 different mammalian cell types (Zheng et al., 2025). To investigate the correlation between IDR self-association and TE, we categorized the IDRs into top and bottom quartiles based on their Epsilon score, a measure of the mean-field inter-residue interaction energy. The bottom quartile represents IDRs with the greatest predicted net self-interaction (Ginell et al., 2025). We found that self-attractive IDRs tend to have a higher TE, but also tend to occur in shorter proteins (**Fig. S3B**). To account for the known anticorrelation of TE with protein length ((Weinberg et al., 2016; Zheng et al., 2025); and recapitulated here in **Fig. S3A**), we then compared the predicted TE of length-matched top and bottom quartile IDRs (by quintiles of total protein length). This analysis revealed a consistent trend: attractive IDRs tend to have lower TE (**Fig. 5A, S3C**). In other words, they are generally translated in a fashion that disfavors self-association.

**Figure 5:**
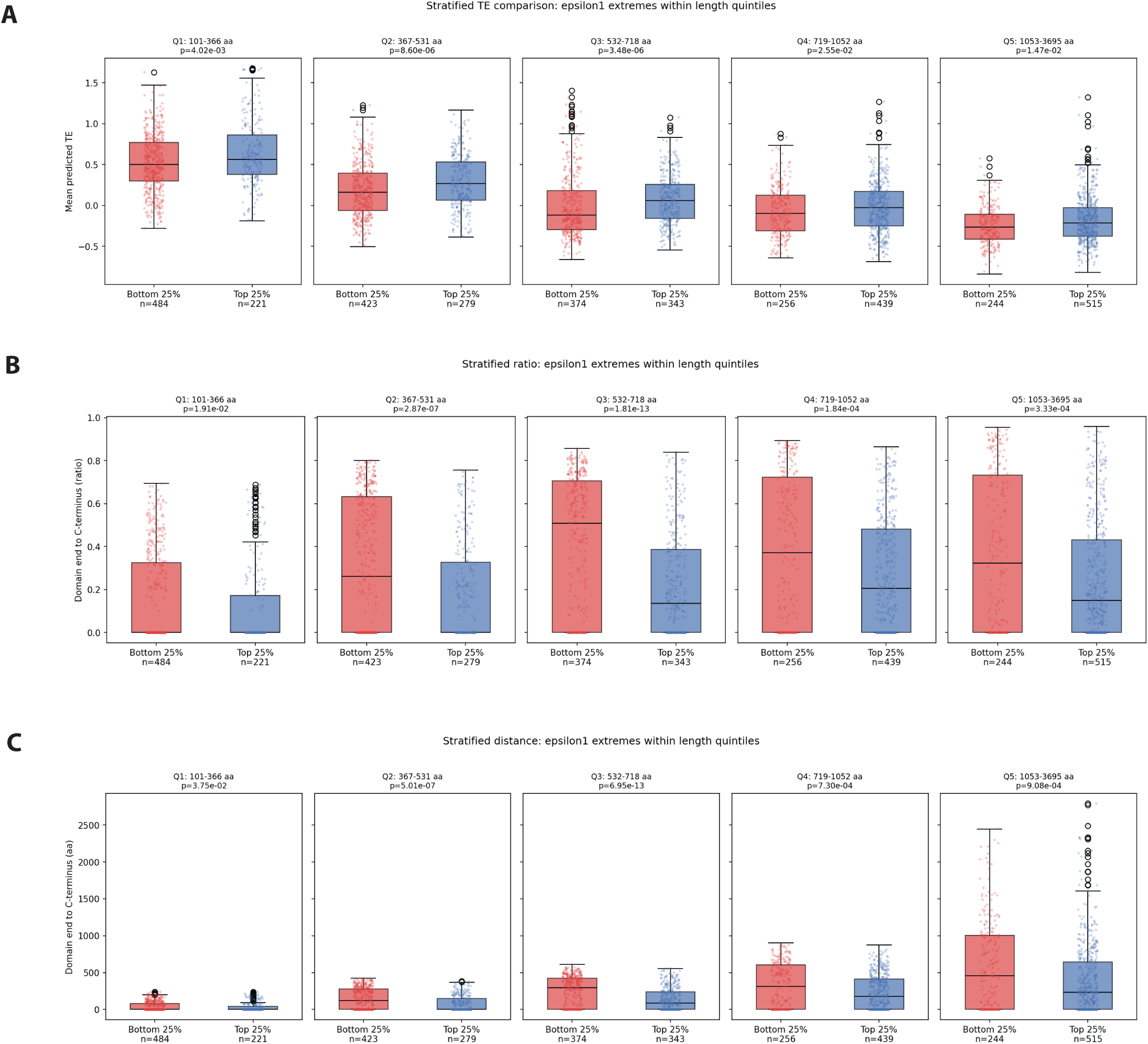
Computational analysis implicating translational control of IDR self-association across the human proteome. (A) Comparison of TE between the attractive (Bottom 25%) and repulsive (Top 25%) IDR containing proteins while accounting for protein length (by categorizing them into quintiles by length) reveals a significant reduction in Mean predicted TE among attractive IDR-containing proteins. Analysis of distance from the end of the IDR to the C-terminal end of the protein shows an unequivocal preference for longer tails among attractive IDRs, be it by (B) raw length in residues or by (C) fraction of total protein length.

Translation efficiency is broadly controlled by the cell, with transcripts that are normally translated poorly often becoming more efficient under stress. To assess whether the self-attractive IDRs are positioned for increased assembly when cells conditionally increase their TE, we evaluated their potential for co-translational retention at polysomes by calculating the coding sequence length from the end of the IDR to the stop codon. We also explored correlation with the distance normalized by total protein length. Strikingly, attractive IDRs tend to have much longer tails (both absolute and relative) than repulsive IDRs (**Fig. 5 B,C, S3 D,E**). We conclude that attractive IDRs in humans are specifically poised for regulation by conditionally accelerated translation.

## Discussion

The prevailing view of age-associated aggregation is that it results from a progressive accumulation of damage (Cuanalo-Contreras et al., 2022; Singh and Kundu, 2026; Tanase et al., 2016). To the extent this view concerns damage to individual protein molecules, it is contradicted by wide-ranging observations that reducing protein biosynthesis prolongs lifespan. The process of replacing old protein molecules with new protein molecules promotes rather than hinders aging.

Our observations shed light on this paradox. We find that reducing the influx of new proteins either directly -- at the level of translation, or indirectly -- at the level of turnover by the proteasome (which in turn reduces translation through negative feedback), sharply reduces the frequency of amyloid aggregation in cells. In sum, our data establish that global equilibration of the supersaturated proteome is promoted by local nonequilibrium: even as translation replaces old protein molecules that may be *thermodynamically* predisposed to aggregate, it creates new molecules that are *kinetically* predisposed to aggregate. The aggregate nuclei formed by those new molecules go on to template the aggregation of old molecules that may exist at globally supersaturated concentrations. Hence, rather than preventing the formation of stable aggregates, translation and the attendant clustering of nascent chains on polysomes accelerates it. More broadly, our findings suggest that the kinetics of protein birth at a single mRNA can be as important as steady-state abundance in determining whether the population of that protein species in the cell remains functional or instead aggregates irreversibly.

However, these pathways are generally viewed as static structural constraints of the ribosome. While co-translational assembly is observed as a localized phenomenon for specific native homo-oligomers (Bertolini et al., 2021; Mallik et al., 2025; Shiber et al., 2018; Wruck et al., 2025), the consequences of translational flux on the kinetic landscape of the supersaturated proteome remain largely unexplored. Our data point to polysomes being fertile ground for nucleating interactions of nascent chains of self-assembling proteins, with flux as well as residence time on the polysome promoting nucleation. This was true, even of nucleation-limited globular DFDs which can be fully self-assembled to equilibrium by simply locally concentrating it (Rodriguez Gama et al., 2026).

This behavior is consistent with theoretical models of co-translational condensation (CTC), where N-terminal “strong-interaction” regions are more effectively exposed on the outer branches of the polysome to facilitate association, whereas C-terminal architectures suppress these interactions because the critical regions are released immediately upon completion (He et al., 2025b). Vectorial emergence from the ribosome, transient confinement on polysomes, and physical entanglement within pre-nucleation clusters may explain how even conformationally limited PrDs remain hypersensitive to local polysome dynamics (Kar et al., 2022; Soto-Santarriaga and Clark, 2025; Wu et al., 2026). We deduce that translational flux and co-translational residence act as potent sources of intracellular disequilibrium, allowing proteins to defy standard mass-action kinetics (Marciano et al., 2022).

This model also redefines our understanding of stress-induced aggregation, which has long been known to occur co-translationally (Zhou et al., 2014). Stress-induced translation shutdown could be a protective mechanism to prevent disorder-rich sequences from finding nucleating conformations at a vulnerable time, that would persist long after the stress recedes. This also has immediate implications for other regulated self-assembly systems, including innate immune signaling.

Inflammasome priming provides a useful parallel. TLR4 stimulation induces a burst of ASC translation from polyribosomes, linking innate immune activation to acute production of a nucleation-prone death-fold adaptor, ASC (Haggadone et al., 2026). In that context, ASC is not simply a passive pool of pre-existing adaptor protein; its translational production is dynamically coupled to the timing of inflammasome assembly. Our data suggest that this bursting not only increases global protein quantity, but also acutely increases the local availability and supersaturation of assembly-competent molecules, ensuring rapid, switch-like polymerization during an immune response (Rodriguez Gama et al., 2026, 2022).

Together, these findings establish that translation is not merely a source of new protein mass, but a spatially organized kinetic process that dictates whether supersaturated proteins remain soluble, assemble productively, or collapse into highly ordered self-assemblies. By placing the precise nature of protein synthesis at the center of nucleation control, this work offers a unified mechanistic explanation for why attenuating translational flux can suppress aggregation and preserve cellular proteostasis across stress, aging, and regulated signaling contexts.

## Materials and Methods

### Plasmid and Strain Construction

Unless specified, all Open Reading Frames (ORFs) were codon-optimized for expression in *S. cerevisiae* and cloned into derivatives of yeast vectors V08 and V12 (Khan et al., 2018). For mammalian studies, ORFs were cloned into pcDNA3.1 derivatives. A□42 was codon optimized for human. Full-length protein sequences and plasmid maps are listed in Table S1.

All yeast strains used in this study are in the S288C background and described in Table S2. Standard DAmFRET experiments were conducted in yeast strains rhy1713, rhy1852, and rhy2145b (WT control for rqc2 experiment) as previously described (Khan et al., 2018; Kimbrough et al., 2025b). rhy1986-1 (integrated with μNS-NLRP3 PYD) was constructed as in (Holliday et al., 2019) by transforming an AseI digest of rhx1950 into rhy1903 to replace the counterselectable *URA3* ORFs. The resulting strain expresses mouse NLRP3 PYD fused to μNS-mTagBFP2 under the control of a doxycycline-repressible promoter. Strain rhy3381 (“rqc2”) was constructed in several steps as follows. rhy1713 was mated with the *pdr5Δ::kanMX* systematic deletion strain and sporulated to create rhy2047, from which rhy2268 was derived by replacing the kanMX marker with natMX from pFA6a-natMX6 (Hentges et al., 2005) by homologous recombination. This strain was then mated with the *rqc2Δ::kanMX* systematic deletion strain, sporulated, and passaged four times on YPD plates containing 3 mM GdHCl (to convert them to [*pin*^−^] (Ferreira et al., 2001)) to create rhy3381.

### Mammalian Cell Culture and Transfection

HEK293T cells were maintained in DMEM supplemented with 10% FBS and 1% Penicillin/Streptomycin at 37°C in 5% CO2. Cells were seeded in 12-well plates and transfected at 60–70% confluence using Lipofectamine 3000 (Invitrogen) according to the manufacturer’s instructions with constructs encoding either mEos-tagged Aβ42 or 3xFLAG-tagged Aβ42. To test the effects of specific pathway inhibitors on amyloid nucleation, cells were allowed to express the transgenes for 24 hours prior to treatment. Cells were then treated with vehicle, Bortezomib (500 nM; R&D Systems, 7282), MG132 (10 µM; VWR, 89161-568), Lactacystin (20 µM; R&D Systems, 2267), Carfilzomib (5 µM; R&D Systems, 7188), Chloroquine (100 µM; Fisher Scientific, AC455240250), or Wortmannin (0.5 µM; MedChemExpress, HY-10197) for 20 hours. Conversely, for HSP70 inhibition experiments, cells were co-treated with BTZ (500 nM) and VER-155008 (20 µM; ApexBio, A4387) immediately upon transfection and maintained for 24 hours. For DAmFRET experiments, cells were fixed using 4% paraformaldehyde (prepared in-house by the institutional core facility) for 5 minutes at 37°C and analyzed following the respective treatment periods.

### Pharmacological Flux Perturbations and SUnSET Assay

To modulate translational flux, HEK293T cells were treated with Bortezomib (BTZ; 1–100 nM) or Harringtonine (25–50 nM; MedChemExpress, HY-N0862) for 20 hours post-transfection. For rescue experiments, media was supplemented with 2 mM Cysteine (Sigma-Aldrich, C7880) to restore translational activity despite proteasome inhibition. In S. cerevisiae, flux was toggled via Rapamycin (200 nM; MedChemExpress, HY-10219), Lactimidomycin (LTM; 3 µM; Sigma-Aldrich, 506291), or Cycloheximide (CHX; 180 nM; Sigma-Aldrich, 01810).Global protein synthesis was quantified using SUnSET (Piecyk et al., 2024); cells were pulsed with 5 µg/mL puromycin (InvivoGen, ant-pr-1) for 15 minutes, and incorporation was detected via Western Blot using an anti-puromycin antibody (Millipore; RRID:AB_2566826).

### Semi-Denaturing Detergent Agarose Gel Electrophoresis (SDD-AGE)

SDD-AGE is a biochemical approach to detect polydisperse assemblies that resist denaturation in ionic detergents (Halfmann and Lindquist, 2008; Kryndushkin et al., 2003), a characteristic of amyloid that derives from its extensive hydrogen bonding network and exquisite side chain packing (Eisenberg and Sawaya, 2017). The SDD-AGE procedure was adapted from (Halfmann and Lindquist, 2008), with the following modifications. Cells were grown to 80% confluency, lysed on ice, and total protein concentration was normalized using a BCA assay. 2% SDS was used in the sample buffer with a 5-minute room-temperature incubation, and 3xFLAG-fused protein distributions were analyzed by capillary transfer to a PVDF membrane. Immunoblotting was performed using a rabbit anti-FLAG primary antibody (Sigma-Aldrich, F7425) and an HRP-linked anti-rabbit secondary antibody (Cell Signaling Technology, 7074). Signals were detected via chemiluminescence (Thermo Fisher Scientific, 34095) and imaged using Image Lab software (Bio-Rad).

### Antibody Generation and Immunoprecipitation

A custom polyclonal anti-mEos antibody was generated and affinity-purified in New Zealand Rabbits against recombinant mEos3.1 protein by GenScript. The resulting affinity-purified antibody (Rabbit #4552) demonstrated a purity of ≥97% by SDS-PAGE and was supplied at a concentration of 0.582 mg/mL. Following the 20-hour treatment period with 500nm BTZ, cells were harvested and lysed. Immunoprecipitation (IP) of the mEos-tagged complexes was performed using the Pierce™ Classic Magnetic IP/Co-IP Kit (Thermo Fisher Scientific, Cat. No. 88804) according to the manufacturer’s instructions, utilizing the custom anti-mEos antibody for capture. For immunoblot validation, fractions were transferred to a PVDF membrane and probed using the custom rabbit anti-mEos primary antibody followed by an HRP-linked anti-rabbit secondary antibody (Cell Signaling Technology, 7074). Signals were detected via chemiluminescence (Thermo Fisher Scientific, 34095) and imaged using Image Lab software (Bio-Rad).

### Mass Spectrometry and Protein Quantification

Eluted protein complexes from the IP and Flow Through fractions were reduced, alkylated, and enzymatically digested with trypsin. The resulting peptides were desalted and analyzed via liquid chromatography-tandem mass spectrometry (LC-MS/MS) using an Evosep One HPLC system (Evosep) operating on the 40-SPD method, coupled to a timsTOF mass spectrometer (Bruker Daltonics). Data was acquired in DIA-PASEF mode using the FleX method settings. Downstream data processing was conducted using label-free quantification (LFQ). Raw MS data were processed using Spectronaut 20 (Biognosys) to identify peptides and quantify protein abundance.

### Bioinformatic and Statistical Analysis of Proteomic Datasets

Statistical analyses of the proteomic count data were conducted to determine relative protein abundance across conditions. Principal Component Analysis (PCA) was performed to assess sample variance and clustering. Differential expression between comparisons (e.g., BTZ-treated vs. vehicle) was assessed to generate log₂ fold changes (log₂FC) and adjusted P-values.

Significantly downregulated proteins were subjected to pathway enrichment analysis using ShinyGO. Query genes were mapped to ENSEMBL gene IDs or STRING-db protein IDs. Enrichment statistical significance was calculated using a hypergeometric test, and raw P-values were adjusted using the Benjamini-Hochberg method to yield False Discovery Rates (FDR). Fold enrichment was calculated as the ratio of the percentage of input genes matching a pathway to the percentage of background genes in that same pathway. The background gene set was defined as the total number of unique genes in the chosen database. Pathways were filtered by an FDR cutoff of < 0.05 and sorted sequentially by FDR and Fold Enrichment.

### Network and Clustering Visualization

To interpret overarching biological themes, significant pathways were clustered based on shared gene overlap. Hierarchical clustering trees were generated to group pathways sharing extensive gene sets, utilizing raw P-values to scale nodes. Pathway interaction networks were constructed by connecting nodes (pathways) that shared ≥20% of their mapped genes, with node size reflecting the total gene count and edge thickness indicating the degree of overlap. Protein-protein interaction (PPI) networks for specific context-dependent protein subsets (e.g., Aβ42- +/- BTZ) were mapped and visualized using STRING-db.

### Accounting for global protein concentration effects of Kozak manipulations of the query protein

Following yeast transformation and selection in -Ura as is typical of our DAmFRET 2-micron expression clones, colonies expressing a strong Kozak sequence were serially passaged in synthetic complete glucose media (with uracil supplementation) up to 4 times on agar petri dishes, to reduce plasmid copy numbers. Likewise, cells expressing a weak Kozak sequence were passaged on agar plates lacking uracil to maintain high plasmid copy numbers. A number of colonies across different passages were inoculated in SD complete media (glucose carbon source) broth, and induced in SGal complete (galactose carbon source) before flow cytometry to assess total protein expression distributions. Colonies with weak Kozak or strong Kozak sequences in passages with overlapping expression distribution were further analyzed via DAmFRET upon mEos photoconversion.

### µNS Seeding Assay

A multivalent viral protein (µNS) was fused to NLRP3 PYD as described previously (Rodriguez Gama et al., 2026) and expressed in trans with Kozak variants of NLRP3 PYD. Because polymerization proceeded to equilibrium, the resulting DAmFRET profiles were analyzed by fitting splines as described previously (Wu et al., 2026).

### frFAST Variant of DAmFRET

Cells were cultured as for a typical DAmFRET experiment and at the harvesting step for cytometry, instead of photoconversion, we added a non-limiting concentration (100 μM) of FRET-compatible dyes tfLime and tfPoppy (Twinkle Factory) and let agitate the cultures for 5 minutes to allow for dye uptake and labeling.

### DAmFRET Data Acquisition

DAmFRET acquisition was conducted in accordance with our previous studies (Khan et al., 2018; Rodriguez Gama et al., 2022). Briefly, individual yeast colonies were grown overnight in SD -ura (2% dextrose) at 30°C and 1000 rpm. Cells were induced in SGal -ura (2% galactose) for 16 hours, followed by resuspension in fresh SGal -ura for 4 hours to minimize autofluorescence. 75 µL of cell suspension was re-arrayed into 96-well plates and photoconverted for 5 minutes using an OmniCure S1000 (320-500 nm filter). HEK293T cells were plated in 12-well plates, fixed with 4% paraformaldehyde for 5 minutes at 37°C and 150 µL of cell suspension was subjected to the same 5 minute photoconversion process. High-throughput flow cytometry was performed on a Bio-Rad ZE5. Donor and FRET signals were collected using a 488 nm laser (100 mW) into 525/35 and 593/52 detectors, respectively. Acceptor signal was collected from a 561 nm laser (50 mW) into a 589/15 detector. Autofluorescence was monitored via a 405 nm laser (100 mW) into a 460/22 detector. Acceptor intensities were normalized by side scatter (SSC) in *S. cerevisiae*, used as a proxy for cell volume, to calculate arbitrary concentration units. Unadjusted acceptor intensity was used as the concentration proxy in HEK293T cells due to morphological variance.

### DAmFRET Analysis and Statistical Evaluation

FCS 3.1 files were gated for single, expressing cells. DAmFRET plots were divided into 64 logarithmically spaced bins and categorized as explained previously (Wu et al., 2026). Namely, a negative control population was used to define the “monomeric” FRET gate (99th percentile). Cells falling above this gate were classified as FRET-positive, and the fraction of cells with self-assembled protein (f_pos_) was plotted against concentration. To evaluate the nucleation profiles, data were analyzed in GraphPad Prism by fitting the raw f_pos_ values to an unconstrained sigmoidal dose-response (variable slope) model. All parameters—including the baseline, maximal assembly plateau, Hill slope, and log EC_50_—were left completely unrestricted to converge freely based on the experimental data points. Statistical significance was determined using the Extra Sum-of-Squares F-test on the non-linear regressions.For bar graph quantifications, the total percentage of amyloid within the sample was calculated by the high-AmFRET population in a concentration bin that populated both low-and high-AmFRET cells for all comparisons, and statistical significance across treatments was determined using unpaired Student’s t-tests/two-way ANOVA. All data represent three independent biological replicates (n=3) and were analyzed using GraphPad Prism. Figures were compiled using Biorender and Adobe Illustrator.

### Molecular diffusion analysis

*In vivo* microrheology was performed using 50 nm Genetically Encoded Multimeric nanoparticles (GEMs) (Delarue et al., 2018; Hernandez et al., 2024). Following transfection with the 50 nm GEM construct and subsequent treatment with either vehicle or BTZ, cells were imaged using high-speed time-lapse fluorescence microscopy. GEM dynamics were captured at a temporal resolution of 10 ms per frame.

Single-particle tracking was employed to extract individual GEM trajectories. For each trajectory, the mean squared displacement (MSD) was calculated to quantify particle mobility. The effective diffusion coefficient (*D_eff_*) was determined by fitting the initial trajectory data to the diffusion equation *MSD = 4D_eff_t.* Fits were specifically restricted to a 100 ms timescale, utilizing the first 10 time points (10 ms intervals) of each tracked movie. To account for intracellular heterogeneity, measurements were performed on robust datasets comprising multiple cells per condition, with hundreds to thousands of individual GEM trajectories captured per cell. For final statistical comparisons and visualizations, *D_eff_* was reported using the median values across datasets, with variance expressed as the standard error of the mean (SEM).

### Relationship of IDRs and translation efficiency

#### Extraction of IDR Interaction Parameters

Intrinsically disordered region (IDR) sequences and their corresponding homotypic interaction parameters (epsilon) were obtained from the FINCHES (Fast IDR proNeness to CoHesion Evaluated from Sequence) dataset generated by (Ginell et al., 2025). For a total of 7,872 unique human proteins (all with IDR >100 residues), we extracted the homotypic epsilon values from the source paper calculated using two distinct coarse-grained force field models: Mpipi-GG and CALVADOS2. Epsilon represents the mean-field interaction energy of the sequence. Negative values indicate attractive homotypic interactions—representing a thermodynamic propensity for self-assembly—whereas positive values indicate repulsive interactions.

#### Protein annotation

Protein lengths and Ensembl gene cross-references were retrieved from UniProt (REST API, batch queries). UniProt IDs were mapped to Ensembl gene IDs via the cross-reference field; only one-to-one mappings were retained. Domain-to-C-terminus distances were computed in absolute (aa) and fractional (of the full length) terms.

#### Translation efficiency prediction

The RiboNN deep learning model was deployed locally with pretrained human multi-task weights. 5′UTR, CDS, and 3′UTR sequences were extracted from Ensembl v110 (GRCh38) for all 89,096 protein-coding transcripts (Zheng et al., 2025). After filtering for valid CDS and length constraints, 65,118 transcripts received TE predictions across 78 cell types; the mean predicted TE was used for downstream analysis.

#### TE assignment

Each protein was assigned the mean predicted TE of its Ensembl canonical transcript (mean of 78 values, each one for a cell-type). Multi-IDR proteins were reduced by keeping the domain with the lowest epsilon (separately for each epsilon score). Proteins without Ensembl mappings, canonical transcripts, or valid predictions were flagged and excluded from downstream analysis (779 total: 506 no Ensembl mapping, 261 excluded from prediction, 11 invalid domain coordinates, 1 no canonical transcript).

#### Downstream analysis

Group comparisons used the Mann–Whitney U test (two-sided) and Kruskal–Wallis test. Correlations were assessed with Pearson r and Spearman ρ. Length-stratified analyses binned proteins into quintiles by protein length, then compared TE between epsilon extreme groups (bottom and top 25%) within each bin.

## Acknowledgements

We thank Brian Slaughter, Paula Berry, Justin Mehojah, Xiaoqing Song and Brooklyn Lerbakken for assistance with experiments and Mark Miller for assistance with illustrations. We thank Ariel Bazzini and Onn Brandman for productive discussions about this work. This work was supported by the American Cancer Society (RSG-19-217-01-CCG to RH), the National Institute of General Medical Sciences (Award Number R01GM130927, to RH) and the National Institute on Aging (Award Number F32AG077876, to AVS) of the National Institutes of Health, and the Stowers Institute for Medical Research. The funders had no role in study design, data collection and analysis, or manuscript preparation. The content is solely the responsibility of the authors and does not necessarily represent the official views of the funders.

## Data availability

Original data underlying this manuscript can be accessed from the Stowers Original Data Repository at http://www.stowers.org/research/publications/libpb-2634

---

**Supplemental Figure 1:**
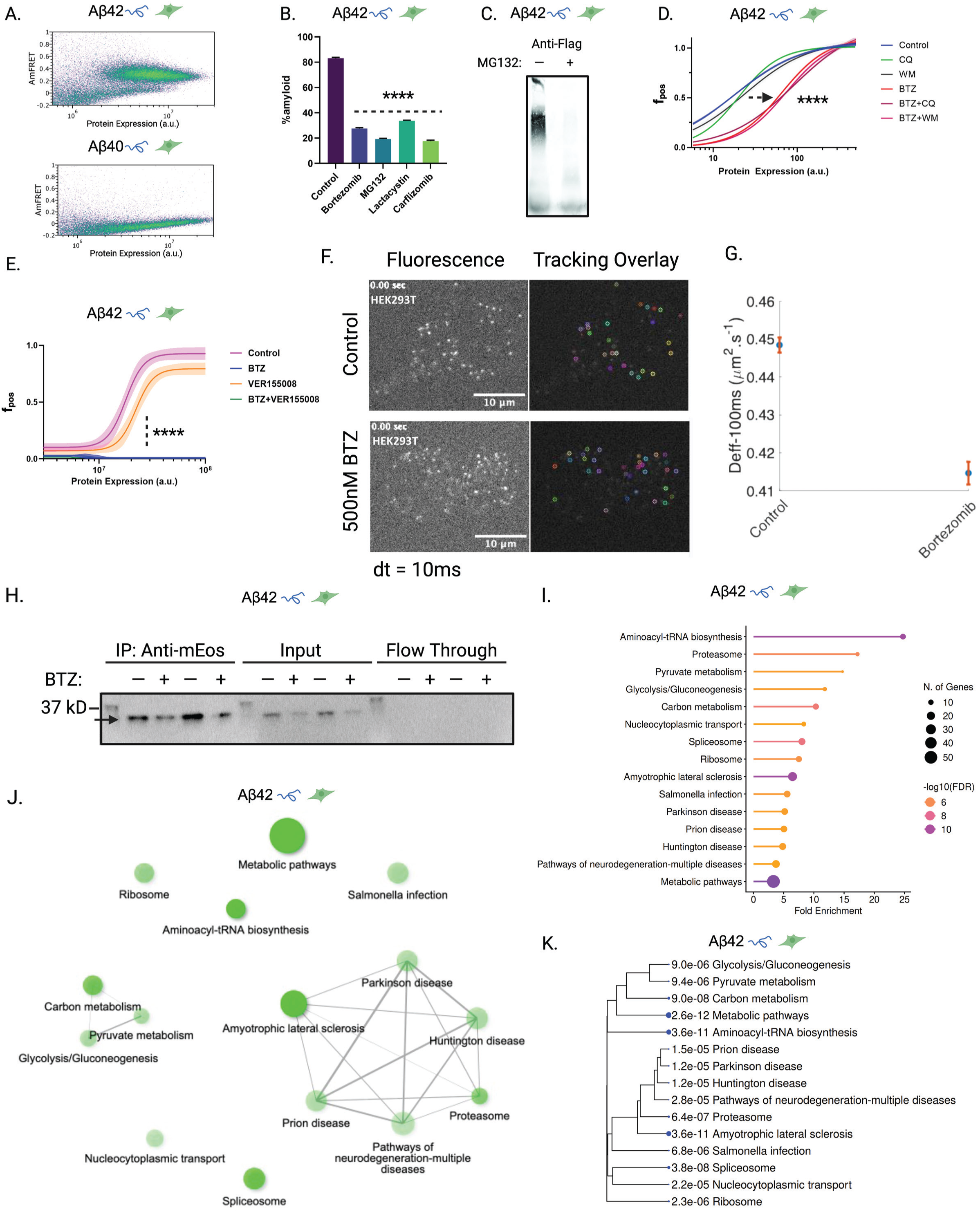
Proteasome inhibition suppresses Aβ42 amyloid nucleation independent of fluorescent tagging, compensatory autophagy, or chaperone upregulation. (A) Representative DAmFRET plot of HEK293T cells expressing mEos-tagged Aβ42 or Aβ40 showing Aβ42 forms a discontinuous DAmFRET profile, characteristic of nucleation-limited phase transitions, including amyloids, while Aβ40 exhibits a single low-AmFRET population. (B) Quantification of amyloid formation via DAmFRET-gating following treatment with diverse classes of proteasome inhibitors. HEK293T cells expressing mEos-Aβ42 were allowed to express the construct for 24 hours prior to a 20-hour treatment with vehicle (Control), 500 nM Bortezomib, 10 µM MG132, 20 µM Lactacystin, or 5 µM Carfilzomib. Broad inhibition of amyloid nucleation across all tested compounds confirms the effect is driven by proteasome impairment rather than an off-target activity of Bortezomib. (C) SDD-AGE analysis of cell lysates expressing 3xFLAG-tagged Aβ42. Cells were allowed to express the construct for 24 hours and were subsequently treated with vehicle (–) or 10 µM MG132 (+) for 20 hours. Anti-FLAG immunoblotting confirms that proteasome inhibition substantially reduces amyloid formation, demonstrating that the observed suppression is not an artifact of the bulky mEos fluorescent tag. (D) DAmFRET non-linear regression fits representing the fraction of assembled Aβ42 as a function of protein expression (a.u.). Following 24 hours of expression, HEK293T cells were treated for 20 hours with vehicle (Control), 500 nM Bortezomib (BTZ), 100 µM chloroquine (CQ), 0.5 µM wortmannin (WM), or co-treated with BTZ+CQ or BTZ+WM. The inhibition of autophagy by CQ or WM failed to rescue the BTZ-induced suppression of nucleation, indicating that enhanced autophagic clearance is not responsible for the increased kinetic barrier to assembly. (E) DAmFRET non-linear regression fits for HEK293T cells treated with vehicle, 500 nM BTZ, 20 µM of the broad-spectrum HSP70 inhibitor VER-155008, or a combination of BTZ and VER-155008. Cells expressed mEos-Aβ42 with drug treatment upon transfection and were measured after 24 hours. Blocking the primary axis of the heat shock response did not rescue the BTZ-induced nucleation defect, suggesting that an upregulation of chaperones is unlikely to explain the loss of assembly competence. (F) Representative raw fluorescence images (left) and corresponding single-particle tracking trajectory overlays (right) of 50 nm Genetically Encoded Multimeric nanoparticles (GEMs) expressed in HEK293T cells. Cells were treated with either a vehicle control or 500 nM Bortezomib (BTZ) for 20 hours. High-speed time-lapse imaging was performed with a temporal resolution of 10 ms per frame (dt = 10 ms). (G) Quantification of the effective diffusion coefficient (D_eff_) evaluated at a 100 ms timescale. Data points represent the median D_eff_ across datasets containing multiple cells and thousands of individual GEM trajectories per condition, with error bars denoting the standard error of the mean (SEM). The slight (<10%) reduction in diffusivity (from ∼0.448 to ∼0.415 µm²/s) upon BTZ treatment indicates that the observed suppression of amyloid nucleation is not driven by a global increase in macromolecular crowding. (H) Immunoprecipitation (IP) validation of mEos-tagged proteins. HEK293T cells were transfected and concurrently treated with vehicle or 500 nM bortezomib (BTZ) for 24 hours. Total cell lysates (Input), IP fractions (Anti-mEos), and depleted lysates (Flow Through) were analyzed by immunoblotting. The arrow denotes the successful enrichment of the ∼26 kD target band across conditions. (I) Kyoto Encyclopedia of Genes and Genomes (KEGG) pathway enrichment analysis of significantly downregulated proteins identified via mass spectrometry following BTZ treatment. The lollipop chart displays top-ranked pathways filtered by adjusted *P* value. The x-axis represents fold enrichment (the percentage of list genes in a pathway divided by the corresponding percentage in the background gene set). Node color indicates statistical significance (-log10(FDR), adjusted via the Benjamini-Hochberg method), and node size corresponds to the number of mapped genes overlapping with the target pathway. (J) Pathway interaction network plot illustrating biological overlap between the enriched KEGG pathways. Nodes represent individual pathways, with node sizes scaled to the total number of genes in the respective gene set. Edges connect nodes that share ≥20% of their mapped genes, highlighting highly interconnected biological networks. (K) Hierarchical clustering tree (dendrogram) summarizing the correlation among significantly enriched pathways. Pathways are clustered based on the proportion of shared genes, revealing overarching biological themes systematically suppressed by BTZ treatment. Raw *P* values, calculated using a hypergeometric test, are annotated adjacent to each pathway. All data represent biological triplicates (*n=3*), with error bars indicating mean ± SEM (*\*p < 0.05, p < 0.01, ***p < 0.001, ****p < 0.0001*).

**Supplementary Figure 2:**
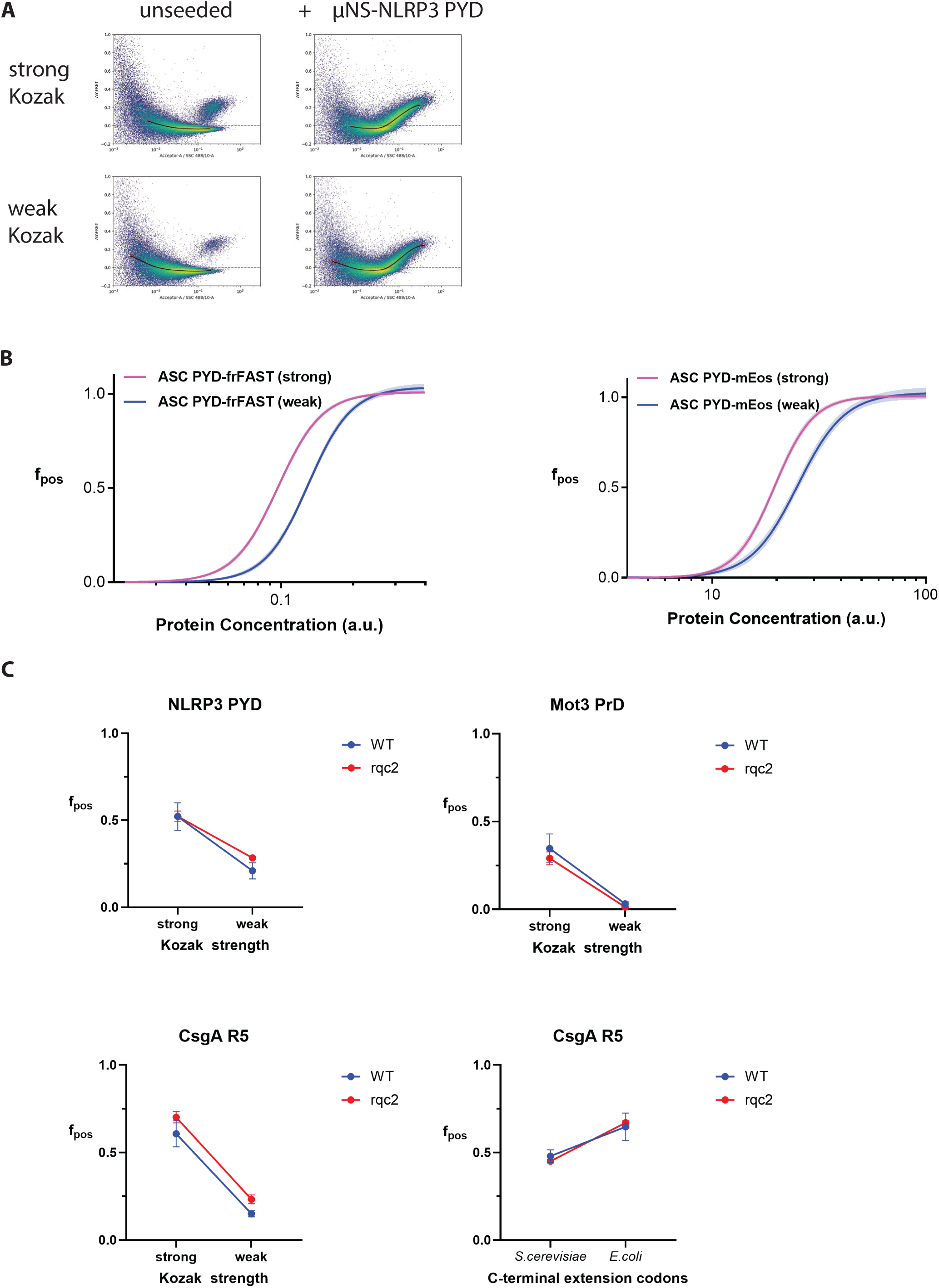
Translation controls kinetic barriers to self-assembly broadly. (A) Kozak perturbation controls the kinetic and not thermodynamic barrier to formation of protein self-assembly. (Left) Montage of representative DAmFRET plots with spline fit overlays from yeast expressing NLRP3 PYD-mEos showing the Kozak dependence of nucleation and (Right) comparable equilibrium phase transition when seeded ectopically with μNS-NLRP3 PYD. (B) Kozak effect of nucleation is independent of the fluorescent tag. DAmFRET fits showing qualitatively similar effect of Kozak strength while tagged with a smaller fluorogenic tag, frFAST, orthogonal to the mEos-DAmFRET setup. (C) Translation-mediated control of nucleation does not depend on co-translational ribosome quality control signaling. Comparison of fraction of nucleated polymers in overlapping expression bins show the persistence of both, the Kozak effect as well as that of codon composition of the C-terminal extension in wild-type (WT) and rqc2 (RQC2 deletion strain) (not significant by Two-Way ANOVA).

**Supplemental Figure 3:**
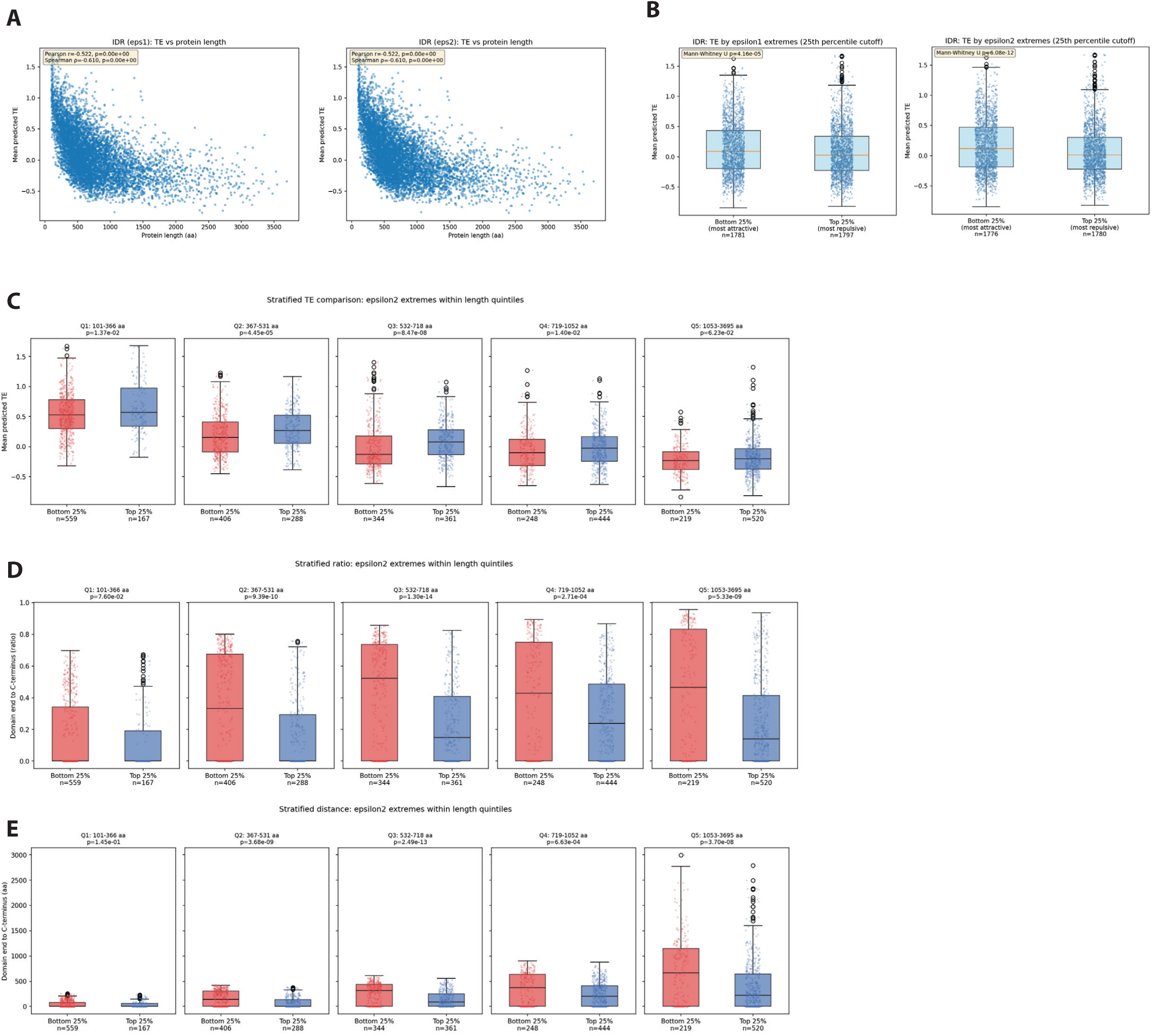
Computational analysis of translation efficiency of IDRs in the human proteome shows co-translation interactions potentially controls IDR behavior. (A) Mean predicted TE via RiboNN showing a characteristic negative dependence on protein length. (B) Comparison of first (Bottom 25%, attractive) and last (Top 25%, repulsive) quartiles by Epsilon score (net self-attractiveness) showing elevated TE for the former, explained by attractive IDRs being present in shorter proteins. (C-E) Data represents the exact content of main Figure 5 A-C, but for the alternate computed self-interaction score (Epsilon2).

